# Negative T cell selection on non-random peptides promotes robust self-nonself discrimination

**DOI:** 10.1101/403428

**Authors:** Inge M. N. Wortel, Can Keşmir, Rob J. de Boer, Judith N. Mandl, Johannes Textor

## Abstract

Our adaptive immune system has the remarkable ability to distinguish previously unseen foreign peptides from harmless self. This self-foreign discrimination was long thought to arise from the silencing of self-reactive T cells during negative selection in the thymus, but recent data show that negative selection is far from complete. Here we ask how a repertoire containing many self-reactive T cells can nevertheless discriminate self from foreign. We address this question using realistic-scale computational models of the T cell repertoire. Our models show that moderate T cell cross-reactivity automatically skews the post-selection repertoire towards peptides that differ systematically from self. But even when no systematic differences between self and foreign exist, discrimination remains possible if the peptides presented in the thymus are chosen in a way that minimizes the co-occurrence of similar, redundant self peptides. Thus, our model predicts that negative selection on a well-chosen subset of self peptides biases the resulting repertoire towards better detection of both self-similar and -dissimilar pathogens. This effect would allow the immune system to “learn self by example”, an ability shared with cognitive systems.

---

To eliminate pathogens without damaging healthy cells, the immune system must discriminate between self and foreign (nonself). The innate arm of the immune system is able to do so with a limited number of germline-encoded receptors that recognize pathogen-associated molecular patterns. By contrast, the adaptive arm of the immune system, which is found in all jawed vertebrates and is mediated by T and B lymphocytes, uses a vastly diverse repertoire of receptors to generate specific protective responses against any pathogen it encounters (1, 2). For example, humans have a repertoire of at least 10^7^ different T cells (3), each expressing one or two of the >10^15^ unique receptor sequences that can arise from the stochastic recombination of V(D)J gene segments and addition of non-templated nucleotides (4, 5). These T cell receptors (TCRs) recognize short foreign peptides presented on major histocompatibility complex (MHC) molecules on the surface of infected or cancerous cells.

However, the random TCR generation process inevitably also produces TCRs that recognize self peptides presented by healthy cells. It was long thought that the majority of these self-reactive receptors are effectively eliminated during T cell development in the thymus through a process termed negative selection (6), but recent studies have shown that this process is nowhere near as complete as it was thought to be (7–9). In fact, given that T cells may only encounter an estimated 10^3^-10^5^ different peptides during negative selection – a small fraction of all MHC-binding self peptides – it is not trivial how negative selection can achieve self-foreign discrimination at all (10–12).

Here, we use computational models to investigate under which conditions negative selection can promote self-foreign discrimination, given that T cells are only exposed to a subset of self peptides. We show that to a certain extent, T cell repertoires can robustly learn “self” from an incomplete set of examples if T cells are moderately cross-reactive, and (2) the subset of self peptides presented in the thymus is not random but chosen in a way that reduces redundance.

## Results

### An artificial immune system discriminates self from foreign after negative selection

To investigate how incomplete negative selection can still foster effective self-foreign discrimination, we devised an “artificial immune system” (AIS) (13). Our AIS is an algorithmic model of a T cell repertoire (14), similar to how an artificial neural network (ANN) is an algorithmic model of the central nervous system. Because it was important to consider T cell repertoires of realistic scale and complexity, we exploited data compression techniques that allow building AISs containing billions of TCRs (15).

Like ANNs, AISs are not only used for *in silico* modelling of the biological system, but also as general-purpose classification algorithms. We took advantage of this property by first using a well-interpretable classification problem outside of immunology to investigate how a TCR repertoire could discriminate a foreign peptide from a self peptide it has not encountered during selection. Specifically, we built an AIS that distinguishes English from other languages based on short strings (letter sequences) of text. This artificial problem mimics the task of self-foreign discrimination because in both cases, classes (languages or proteomes) are to be distinguished based on a limited amount of information (short strings or peptides). A useful property of the language problem is that it can take on a range of difficulties, as very dissimilar languages such as English and the South-African language Xhosa are much easier to distinguish than related languages such as modern and medieval English.

Our model belongs to the family of “string-based” AISs (10, 14–16) that represents each TCR as a binding motif, and defines a TCR’s affinity for a string as the maximum number of adjacent positions where this motif matches the string (Fig. 1A) (Methods in SI Appendix). A TCR is defined to *react* to all strings for which it has an affinity of at least some threshold *t*, which represents a functional response threshold rather than a mere binding threshold. Crucially, reaction does not require a perfect match between the string and TCR motif. Thus, our TCRs are *cross-reactive* and react to multiple, related peptides. In contrast to models based on binding energy (17, 18), the “motif-based” recognition implemented in our model (Fig. 1A) ensures that both peptides recognized by the same TCR and TCRs recognizing the same peptide share sequence motifs – in line with observations from TCR-specific peptide sets (19–21) and peptide-specific TCR repertoires (22, 23).

**Fig. 1.**
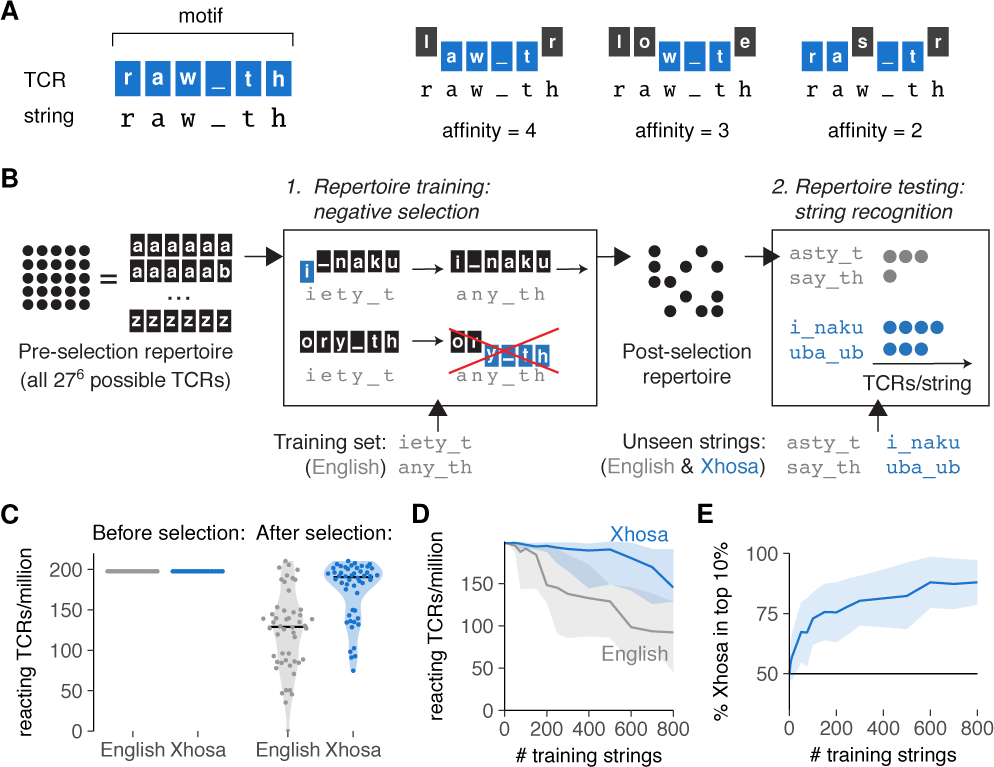
Negative selection on a subset of the whole “self” can achieve self-foreign discrimination. (a) Our model of string recognition represents TCRs by a *binding motif* – the string they bind to most strongly (left). Their affinity for any given string equals the maximum number of adjacent positions where the binding motif matches the string (right). (b) Simulating negative selection *in silico*: (1) TCRs in the unbiased pre-selection repertoire (with all possible 27^6^ ≈ 400 million TCR motifs of 6 characters [a-z and _]) are deleted if their affinity for any of the *training strings* exceeds the functional response threshold *t*. (2) Unseen English and Xhosa strings are exposed to the post-selection repertoire to find the number of remaining TCRs reacting to them (that is, TCRs with affinity ≥ *t)*. (c) Reacting TCRs per million of unseen English and Xhosa strings, before and after negative selection on 500 English strings. Horizontal lines indicate medians. (d) Median and interquartile range of English-and Xhosa-reactivity after negative selection on English strings. (e) Percentage of Xhosa strings among the 10% of strings with the most reacting TCRs after negative selection on English strings (mean ± standard deviation, SD, of 30 simulations). No discrimination should result in equal amounts (50%) of English and Xhosa strings in this top 10%. Throughout this figure, we tested 50 English and 50 Xhosa strings using an affinity threshold *t* = 3 for negative selection.

To test how well TCR repertoires could discriminate between two very dissimilar languages (English and Xhosa) after incomplete negative selection, we started with an unbiased pre-selection repertoire with equal numbers of TCRs reacting to English and Xhosa, and then performed *in silico* negative selection on an English *training set* by deleting all TCRs reacting to any of the (<1000) training strings (Fig. 1B, using a threshold *t* = 3 leading to intermediate cross-reactivity). Although this negative selection did not completely abrogate TCR reactivity towards English strings outside of the training set, it still biased the post-selection repertoire to contain more TCRs reacting to Xhosa than to English (Fig. 1C,D).

Given that peptides to which many TCRs react tend to elicit stronger immune responses (24), it is important that these most frequently recognized peptides are predominantly foreign. The 10% most frequently recognized strings in our simulation were indeed predominantly Xhosa strings (Fig. 1E). The affinity distribution of these TCR interactions was shifted towards higher affinities for Xhosa, but only very slightly (Fig. S1A). For sake of simplicity, we therefore focus only on the number of reacting TCRs throughout this paper, rather than considering different affinities separately. This choice to consider TCRs with a broad range of affinities is supported by growing evidence that also lower affinity TCRs are important contributors to immune responses (25).

### Discrimination success relies on moderate cross-reactivity and sequence dissimilarity

These results confirm that our AIS can easily distinguish English from Xhosa even after incomplete negative selection. To investigate in more detail under which conditions this discrimination arises, we analyzed which TCRs were deleted during negative selection on English strings (Fig. 2). TCRs reacting to “unseen” English strings (those absent from the training set TCRs were exposed to during negative selection) had a reduced survival compared to TCRs reacting to Xhosa strings (Fig. 2A). Because TCRs are only deleted when they react to at least one string in the training set, this implies that strings eliciting reactions from the same TCRs tend to represent the same language. To visualize this, we created graphs in which each node represents a string, and two nodes become connected neighbors when at least 5 TCRs per million pre-selection TCRs react to both of them (Fig. 2B). Indeed, neighbor strings are largely from the same language (Fig. 2B, left), which is quantified by the *concordance*, the average proportion of neighbors from the same language. To show that the high concordance (0.81) of English and Xhosa strings represents intrinsic differences between English and Xhosa strings, we randomly divided English strings into two groups and constructed a similar graph, which as expected has a concordance of only 0.5 (Fig. 2B, right). This confirms that our TCRs can only discriminate between two sets of strings that are intrinsically different.

**Fig. 2.**
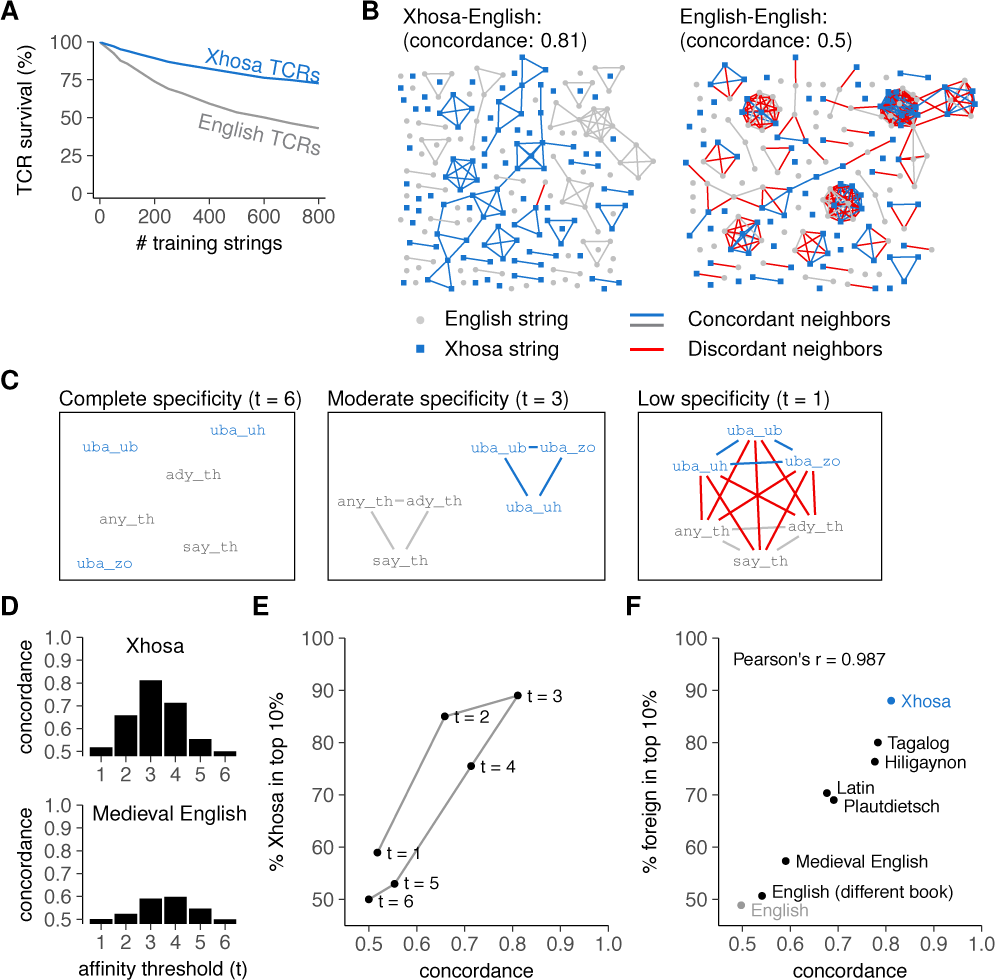
Discrimination requires moderate TCR cross-reactivity and dissimilar self-and foreign strings. (a) Mean percentage of surviving TCRs reacting to English and Xhosa strings after negative selection (using threshold *t* = 3). Plot represents a different analysis of data shown in Fig. 1D,E. (b) String similarity visualized in a graph where nodes (strings) are neighbors (connected by edges) if at least 5/million pre-selection TCRs react to both. (c) Cross-reactivity increases the number of edges between example English and Xhosa strings (demonstrated here for a few examples). Edges between strings from different languages are shown in red. (d) Concordance in the English-Xhosa and English-Medieval English graphs for different thresholds *t*. (e) Concordance and discrimination between English and Xhosa for different thresholds *t*. Negative selection was performed on 800 English strings. Datapoint for *t* = 3 corresponds to the endpoint of Fig. 1E. (f) Language concordance versus enrichment of foreign strings among the top 10% most frequently recognized strings after negative selection (*t* = 3, selection on 800 English strings). Pearson’s correlation coefficient r = 0.977, with 95% confidence interval [0.890, 0.995]. The control “English” compares two sets of English strings from the same book that was used for training (Moby Dick), whereas “English (different book)” compares unseen English strings from the training book to those from the Bible. The point “Xhosa” corresponds to the point “*t* = 3” in Fig. 2E. See also Fig. S1.

Our results indicate two key requirements for achieving self-foreign discrimination through negative selection on an incomplete subset of self: an appropriate level of TCR *cross-reactivity* towards multiple, related strings, and sufficient *dissimilarity* between self-and foreign.

To illustrate the importance of cross-reactivity, we set the affinity threshold in our model to *t* = 6, so that each TCR was maximally specific and only reacted to the one string matching its binding motif perfectly (i.e., no cross-reactivity). The corresponding graph contains no neighbors at all (Fig. 2C, left) and has a concordance of 0.5 (Fig. 2D,E). Consequently, maximal TCR specificity abolishes self-foreign discrimination in our model (Fig. 2E) because without cross-reactivity, negative selection cannot delete TCRs for strings that are not part of the training set – it therefore deletes very few TCRs (Fig. S1B). However, very low specificity (*t* = 1) is equally problematic as it results in a graph where any two strings are neighbors irrespective of language (Fig. 2C, right), which leads to low concordance even between dissimilar languages (Fig. 2D,E), poor self-foreign discrimination (Fig. 2E), and often even deletion of the entire repertoire (Fig. S1B). Only intermediate specificities allow TCRs to preferentially react to either English or Xhosa strings (Fig. 2C, middle). This results in both a high concordance (Fig. 2D,E) and a preference for Xhosa-reactivity in the post-selection repertoire (Fig. 2E).

As shown in Fig. 2B, even an optimal level of cross-reactivity will not result in a high concordance unless the languages are intrinsically different. The accomplished level of self-foreign discrimination therefore depends directly on the similarity between self-and foreign sequences. Indeed, when we repeated our analysis for a number of other languages with varying similarity to English, we found a linear correlation between concordance and the acquired level of discrimination (Fig. 2F). This was a property of the tested languages rather than the specific texts chosen, as our model could not discriminate between English strings from different books (Fig. 2F).

### Sequence similarity hampers discrimination between self-and foreign peptides

These results on natural languages suggest that TCR cross-reactivity and sequence dissimilarity should also be important for self-foreign discrimination in the immune system. We therefore applied our AIS model to self-foreign discrimination by CD8^+^ T cells, which recognize peptides bound to the MHC class I (MHC-I) complex with a typical length of nine amino acids (AAs). The six residues at positions 3-8 are thought to be most relevant for TCR binding (26). Accordingly, we modified our TCR model to accommodate 6-mer peptide sequences rather than six-letter strings (Fig. 3A). Setting the affinity threshold to an intermediate value of *t* = 4 in this model allowed each TCR to react to roughly one in every 55,000 peptides (Fig. S2A) – a cross-reactivity level that reasonably matches an experimental estimate of one in 30,000 (27). Furthermore, at this level of cross-reactivity, peptides elicited reactions from 0 to 20 TCRs per million in our simulated repertoires (Fig. S2B), in line with experimental data (28–31). These results suggest that the cross-reactivity level of TCRs roughly matches that of our model at *t* = 4, well within the “moderate” range allowing discrimination between dissimilar strings (Fig. 2D,E).

**Fig. 3.**
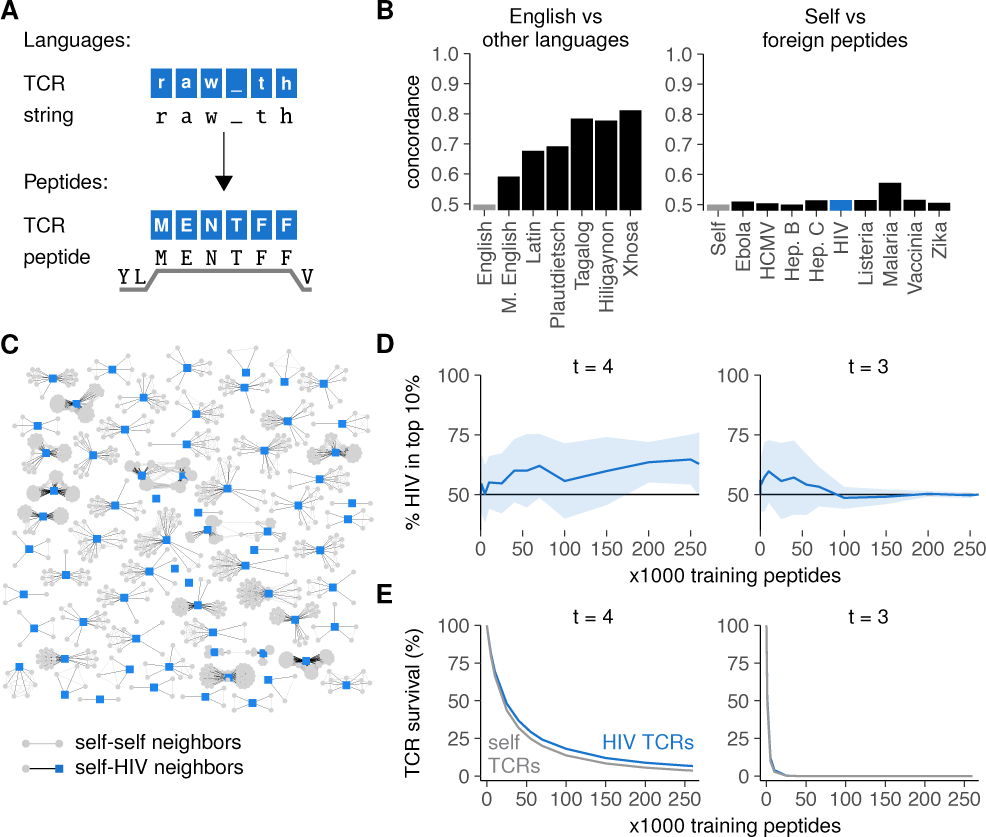
High similarity between self-and foreign peptides hampers their discrimination by the immune system. (a) TCR binding to peptides on MHC-I (HLA-A2:01) focuses on the 6 residues at positions 3-8 and resembles the TCR-string model as in Fig. 1A. (b) Concordance for English versus other languages (left) compared to that for self versus foreign peptides (right). Language concordances from Fig. 2F are included for comparison. (c) Graph of HIV peptides and their neighbors. Edges connect peptides that have at least 5/million preselection TCRs in common. (d) Percentage of HIV-peptides among the 10% most frequently recognized peptides after negative selection (mean ± SD of 30 simulations). (e) Mean percentage surviving TCRs for self and HIV peptides after negative selection.

To examine whether self-and foreign peptides are dissimilar enough to allow self-foreign discrimination, we first predicted MHC-I-binding peptides from the human proteome (32) and used the residues 3-8 as MHC-bound self peptides in our model. To obtain foreign sequences, we predicted MHC binders for a variety of pathogens associated with T cell immunity: the malaria parasite, the bacterium *Listeria monocytogenes*, and the viruses ebola, hepatitis B, hepatitis C, human cytomegalovirus (HCMV), human immunodeficiency virus (HIV), and vaccinia (Table S1).

Graphs of self versus foreign peptides had strikingly low concordances (Fig. 3B)(Methods in SI Appendix), barely exceeding the control concordance observed between two random, different sets of self peptides (“Self", negative control), and lower than the concordance we had observed between modern and medieval English. This was a property of the sequences themselves rather than the chosen threshold *t* (Fig. S3A). In a graph of all HIV peptides and their neighbors, the majority of HIV peptides had many self neighbors whereas none of them had HIV neighbors (Fig. 3C) – indicating that most HIV peptides are more similar to peptides from the human proteome than to other HIV peptides.

This high similarity between self-and foreign peptides suggests that achieving self-foreign discrimination via negative selection is difficult. Indeed, although the realistic cross-reactivity at *t* = 4 allowed some discrimination between self-and HIV peptides as shown by a small enrichment of HIV among most frequently recognized peptides (Fig. 3D, left), this effect came nowhere close to that observed for languages (Fig. 1E), even with very large numbers of training self peptides. Consistent with this observation, the survival of self-reactive TCRs was only slightly lower than that of HIV-reactive TCRs (Fig. 3E, left). These results were not specific for HIV peptides, as we obtained similarly low levels of self-foreign discrimination for all other pathogens tested (Fig. S3B). Self-HIV discrimination was even worse for *t* = 3 and rapidly disappeared completely as TCR survival diminished for large training sets (Fig. 3D,E, right), confirming that self-foreign discrimination becomes more difficult when TCRs are too cross-reactive.

### Selection on non-random peptides greatly improves self– foreign discrimination

Thus, although incomplete negative selection can achieve self-foreign discrimination in principle, achieving sufficient discrimination is very difficult in practice because self-and foreign peptides can be extremely similar and therefore can be recognized by the same TCRs. Clearly, the immune system must overcome this problem in order to balance the removal of self-reactivity with the preservation of foreign recognition. It has previously been suggested that thymic selection should occur on a non-random set of self peptides to achieve self-foreign discrimination (12). We therefore used our model to investigate what an “optimal” set of self peptides would look like, and how much this might improve self-foreign discrimination.

As a starting point, we based the optimization of the training set on the peptide cluster structure as observed in Fig. 3C. The large clusters in this graph contain many similar self peptides, which can delete the same TCRs during negative selection (Fig. 4A). Exchanging one such peptide for one of its neighbors during selection thus has little effect on the post-selection repertoire – and presenting both has little added value. By contrast, self peptides in smaller clusters are far less *exchangeable* (Fig. 4A): their TCRs cannot be removed as easily by other peptides. Thus, negative selection on randomly chosen training sets is inefficient: these sets often contain several exchangeable peptides that delete the same TCRs, while simultaneously missing many non-exchangeable peptides and allowing the corresponding self-reactive TCRs to escape. We therefore used combinatorial optimization techniques (Methods in SI Appendix) to compute peptide combinations that deleted as many different self-reactive TCRs as possible (“optimal” training sets, Fig. 4B). As expected, these optimal training sets contained fewer exchangeable peptides (Fig. 4C, where exchangeability equals the number of self neighbors plus one).

**Fig. 4.**
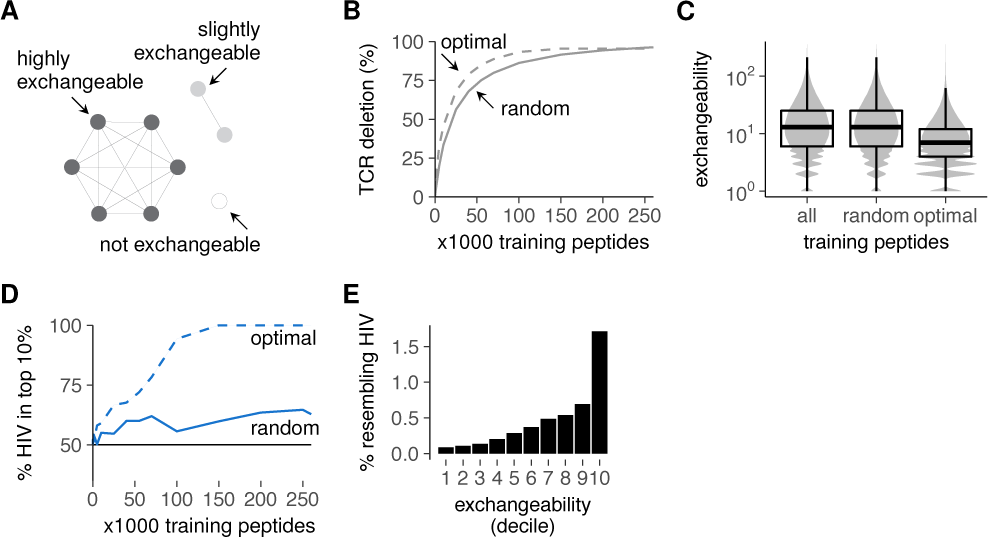
Improved self representation during negative selection allows self-foreign discrimination. (a) Self peptides from large clusters delete the same TCRs as their neighbors and are thus exchangeable during negative selection, whereas peptides from small clusters are not. (b) Mean percentage of self-reactive TCRs deleted by optimal training sets of self peptides during negative selection. TCR deletion with random training sets was computed on the data from Fig. 3E for comparison. (c) Peptide exchangeability distribution in the full set of all self peptides compared to that in random and optimal subsets of 100,000 peptides. Exchangeability is defined as the number of self neighbors + 1. (d) Self-HIV discrimination after selection on optimal training sets. Discrimination after selection on random training sets (Fig. 3D) is shown for comparison. See also Fig. S4. (e) Percentage of self peptides with HIV neighbor(s) plotted against exchangeability (self peptides were divided into 10 equal-number deciles from low to high exchangeability). Negative selection in panels b and d was performed with *t* = 4, and results were plotted as mean±SD of 30 simulations.

We then tested whether these training sets optimized for inducing *tolerance* could also establish self-foreign *discrimination*. This is not guaranteed, as the latter requires not only the removal of self-reactive TCRs, but also the preservation of foreign-reactivity. Nevertheless, our optimal training sets substantially improved self-foreign discrimination (Fig. 4D). This seems to be a consequence of the enrichment for low exchangeability peptides (Fig. 4C), which are less likely to delete HIV-reactive TCRs (Fig. 4E). Importantly, this discrimination still required appropriate TCR cross-reactivity and was absent at *t* = 3 (Fig. S4). From these results, we conclude that negative selection on a representative set of self peptides can alleviate the problem of self-foreign similarity, but only when TCRs are sufficiently specific.

Obviously, our optimal training sets are artificial, and biological negative selection cannot calculate which self peptides should be present in the thymus. We therefore investigated how a representative set of self peptides might reasonably be obtained during real negative selection. Analysis of our optimal training sets revealed an enrichment for rare AAs compared to the total set of self peptides (Fig. S5). Interestingly, peptides with many rare AAs were typically less exchangeable (Fig. 5A). This finding suggests that training sets enriched for rare AAs – similar to our optimal sets – contain fewer exchangeable peptides, and might thus result in better self-foreign discrimination.

**Fig. 5.**
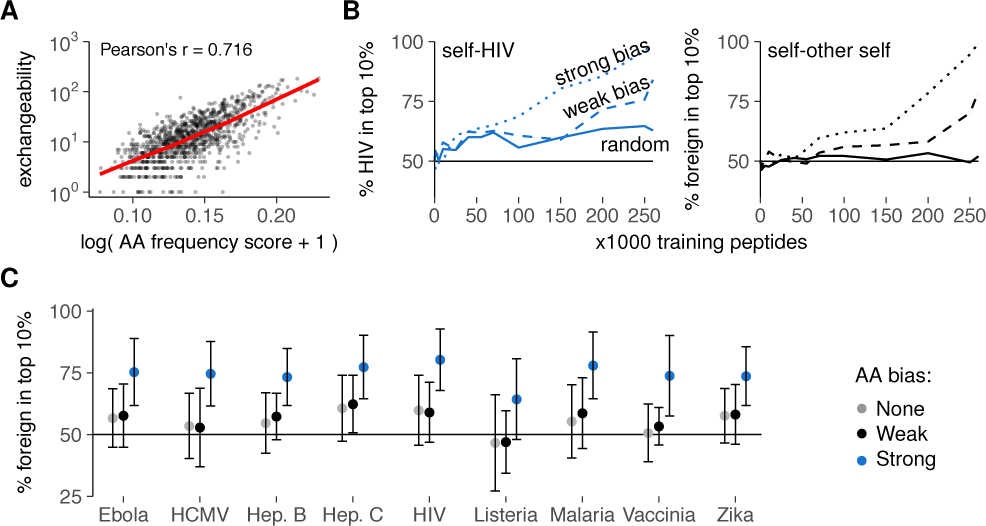
Thymic enrichment for rare AAs facilitates self-foreign discrimination by improving self representation during negative selection. (a) Exchangeability versus peptide AA frequency score in a random sample of 1000 self peptides (frequency score is low for peptides with many rare AAs, (Methods in SI Appendix)). Pearson’s correlation coefficient r = 0.716, with 95% confidence interval [0.684, 0.745]. See also Fig. S5. (b) Discrimination after negative selection on self peptides chosen with a (weak/strong) bias for rare AAs. Discrimination after selection on random peptides (Fig. 3D) is included for comparison. Plots show self-HIV discrimination (left), and self-other self discrimination (right, where a random sample of self was assigned the label “foreign” before selection on training sets from the remaining “self” peptides). (c) Self-foreign discrimination for different pathogens after negative selection on 150,000 self peptides chosen randomly or with AA bias. See Fig. S6 for the full discrimination curves. Negative selection in panels b and c was performed with *t* = 4, and results were plotted as mean±SD of 30 simulations.

To test this hypothesis, we again generated training sets of different sizes, but this time picked our training peptides with a probability that depended on the AA composition of each peptide (Methods in SI Appendix). These probabilities introduced either a weak or a strong bias for self peptides with rare AAs, mimicking the AA enrichment pattern observed in our optimal training sets. This AA bias substantially improved self-foreign discrimination after negative selection, for HIV (Fig. 5B, left) and all other pathogens tested (Fig. 5C, S6). Interestingly, this strategy also worked when we first set aside a random sample of other self peptides as “foreign” before selecting training sets from the remaining “self” peptides. In this scenario, biased training sets still yielded substantial self-“foreign” discrimination, whereas random sets did not (Fig. 5B, right). This result demonstrates that negative selection on non-random training peptides facilitates self-foreign discrimination – even in the extreme case where no inherent difference between self and foreign peptides exists.

## Discussion

Our AIS model explains how negative selection on an incomplete set of self peptides can nonetheless bias a T cell repertoire towards foreign recognition. We demonstrate that a non-random subset of self peptides enriched for rare AAs can balance the removal of self-reactive TCRs with the preservation of foreign-reactive receptors. Importantly, this strategy works even when self and foreign peptides are not inherently different. In fact, for the pathogens we considered, the similarity to self was so high that it is hard to conceive how any self-foreign discrimination could be achieved through negative selection on random peptides. By contrast, a “smart” peptide presentation strategy could still ensure that the peptides best recognized by the immune system are predominantly foreign – even in this difficult scenario. This notion reconciles textbook negative selection theory with recent observations that T cells see only a fraction of all self peptides during thymic selection, and that even healthy individuals have many self-reactive T cells (7).

Although we demonstrate here how negative selection can skew a developing repertoire away from recognition of self, our results also point out that this “central tolerance” alone is likely insufficient for reliable self-foreign discrimination. This is in line with the consensus that peripheral tolerance mechanisms are crucial to prevent and dampen immune responses by those self-reactive cells surviving negative selection. Nevertheless – under the right conditions – negative selection can at least provide a *basis* for such other mechanisms to build on. The idea of a “leaky” central tolerance strengthened by peripheral mechanisms is not new (7, 33), and is supported for example by studies showing that more nuanced discrimination becomes possible when T cells make decisions cooperatively (34, 35). However, our results clearly show that it is not trivial for negative selection to provide even a starting point for self-foreign discrimination. To do so, it must somehow overcome the fundamental problem of similarity between self-and foreign peptides.

Our finding that non-random peptide presentation is a prerequisite for efficient self-foreign discrimination raises the question how the thymus might obtain a preference for presenting low-exchangeability peptides. Although it remains unclear exactly which and how many peptides a T cell sees during selection, the importance of the thymic peptidome in shaping the TCR repertoire is evident from the existence of specialized antigen presenting cells, transcription factors such as AIRE, and even special proteasomes controlling thymic peptide presentation (36). We suggest that the biased presentation of low-exchangeability peptides required for self-foreign discrimination might arise from special binding preferences of thymic antigen presentation proteins. As has already been shown for the thymoproteasome during thymic positive selection (37, 38), such binding preferences can enrich for specific subsets of self peptides and thereby impact the ability of a TCR repertoire to recognize self and foreign. While a bias for specific AAs such as described in this paper would be one way to enrich for low-exchangeability peptides, we do not exclude that other binding preferences could have a similar impact on self-foreign discrimination.

Notably, our imperfect selection accomplishes self-foreign discrimination by also reducing the recognition of peptides the T cell repertoire has not seen during selection. This capability of the T cell repertoire to generalize beyond given examples is a fundamental property of learning systems (39), and allows the repertoire to perform a cognitive task: learning to distinguish self from foreign. Even though this learning process mechanistically differs from learning by the central nervous system, its high-level outcome is remarkably similar, and shares many properties with “slow learning” systems as described in psychology and neuroscience (40).

## Materials and Methods

### Data and code availability

All code used in this paper will be made available at: www.github.com/ingewortel/negative-selection-2018. Data will be made available on www.osf.io.

### Simulation of negative selection

Our general simulation setup can be outlined as follows:

1. Generation of an *unbiased* TCR repertoire containing all possible motifs of length 6. For details, see *Repertoire model of negative selection* (Methods in SI Appendix).
2. Selection of a *training set* of either *n* English strings or *n* self peptides. See *Sequences* for details on the sequences used, and *Training set selection* for details on the manners in which training sets are sampled (Methods in SI Appendix). The training set selection method was random unless mentioned otherwise in the figure legend. The value of *n* can also be found in the figure legend.
3. Negative selection of TCRs on the training set. All TCR motifs that match *any* of the training sequences in at least *t* adjacent positions are removed from the repertoire. Unless mentioned otherwise, negative selection was performed with an affinity threshold *t* = 3 for strings and *t* = 4 for peptides (see figure legends). All TCRs that remain make up the *post-selection repertoire*. For details on computational methods, see *Repertoire model of negative selection* (Methods in SI Appendix).
4. Analysis of the recognition of *test sequences* by the post-selection repertoire. Test sets always consist of “unseen” sequences that were not part of the training set used for negative selection. See figure legends for details on the number and source of the test sequences used. See *Post-selection repertoire analysis* (Methods in SI Appendix) for details on specific analysis metrics used.

We repeat steps 2-4 with different training and test sets for each simulation. In the case of “optimal” training sets, which are per definition selected only in one way (see *Training set selection* (Methods in SI Appendix) for details), the training set was constant across simulations but the test set was varied. Negative selection success as determined by these simulations is then assessed in the context of expectations based on the similarity between self and foreign sequences (see *Sequence analysis* (Methods in SI Appendix) for details).

### Supporting Methods

Detailed computational methods used in this article are available as Supporting Information in the SI Appendix.

## Supporting Information (SI)

The SI Appendix contains Supporting Methods, Figs. S1 to S6, and Table S1.

## Acknowledgements

IW was supported by a Radboudumc PhD grant. JT was supported by a Young Investigator Grant (10620) from KWF. CK and JT were supported by an NWO-ALW grant (823.02.014), and CK was supported by the EU HORIZON2020 program (APERIM project).

## Supplementary Information

### Supporting Information Text

#### Supporting Methods

##### Sequences

We applied our TCR model to both 6-letter strings and 6-AA peptides. Throughout this methods section, we will refer to them as *strings* and *peptides* for methods specific to either languages or peptides, or as *sequences* for methods applying to both. With *self* sequences we mean either human peptides or English strings, and with *foreign* sequences we mean either pathogenic peptides or strings from other languages (see below).

##### Strings

English training strings (“self”) were extracted from Moby Dick (downloaded from **www.gutenberg.org/files/2489/2489.txt**). Independent sets of test strings were extracted from translations of the Gospel of John in the Bible (downloaded from **www.biblegateway.com**). We obtained translations in different languages: English, Medieval English, Latin, and Plautdietsch (Indo-European languages), Tagalog and Hiligaynon (Austronesian languages), and Xhosa (Niger-Congo family of languages). Recognition of these test strings was always compared to recognition of unseen English control strings from the Moby Dick training set. Capital letters were removed and all spaces and punctuation marks were replaced by an underscore (_), yielding text with 27 possible characters (26 letters of the latin alphabet and _). Texts were then randomly cut into strings containing 6 characters each. Please refer to our code repository (see *Data and code availability* in main text) to obtain the exact input text files and the scripts that generate the chunks.

##### Peptides

Proteomes were obtained from Uniprot (1, 2) (Table S1). Potential HLA-A2:01 binders were predicted using NetMHCPan (3) (version 3.0), focusing on peptides of 9 AAs. Using the NetMHCPan default settings, the 2% highest scoring 9-mers were defined as MHC-I binders. Of these, we selected the 6 residues at positions 3-8 to get the TCR-binding 6-mers, and then removed duplicates to get unique 6-mers for each proteome (Table S1).

##### Repertoire model of negative selection

A limiting factor for simulating negative selection on large TCR repertoires is computational complexity. Our unbiased pre-selection repertoires contain TCRs for every possible binding motif of 6 letters (a-z or _) or 6 AAs – resulting in 27^6^ ≈400 million TCRs for the language AIS, and 20^6^ = 64 million TCRs for the peptide AIS. Each of these TCRs needs to be compared against all sequences in the training set. Our implementation of the contiguous affinity model uses advanced computational methods as described in (4, 5) to compress T cell repertoires and to enable these comparisons between large sets of sequences. These methods are available in our code repository (see *Data and code availability* in main text).

##### Training set selection

Training sets of *n* English strings were sampled randomly in each simulation. Training sets of *n* self peptides were sampled from the total ∼260,000 human MHC-I binders in one of three ways: random, optimal, or biased sampling (see below for the last two).

##### Optimal training peptide selection

“Optimal” training sets were designed to remove as many self-reactive TCRs as possible. We listed all self-reactive TCR binding motifs that would react to at least one of the ∼260,000 human MHC-I binders for a given threshold *t*, and then selected combinations of minimal numbers of self peptides that would delete a maximal number of these self-reactive TCR motifs. We could not find an exact solution to this combinatorial optimization problem, because there is a nearly infinite number of ways to select *n* out of ∼260,000 self peptides – and it is not possible to assess the removal of self-reactive TCRs for each of them. We therefore designed a “greedy” algorithm to find an approximative solution instead. Briefly, we iteratively select the self peptides that remove the most remaining self-reactive TCRs by repeating two steps:

1. List the self-reactive TCR motifs that still remain in the repertoire;
2. Select the self peptide that deletes the most of these remaining self-reactive TCRs. If multiple self peptides delete an equal number of remaining TCRs, we pick only those self peptides that do not overlap in the TCRs they delete.

We stop when all self-reactive TCRs are deleted. The result is an ordered list of self peptides, of which the top *n* epitopes form an “optimal” training set of size *n*. For *t* = 3, an optimally chosen 12,025 self peptides (∼5% of all self peptides) could already remove all self-reactive TCRs, whereas this required 130,407 self peptides (∼50% of all self peptides) at *t* = 4. For simulations with optimal training sets larger than this number, random self peptides were added to the optimal combinations to obtain the desired total number *n*.

#### Biased training peptide selection

To generate training sets biased for rare AAs, all self peptides were first assigned a score that depended on their AA composition:

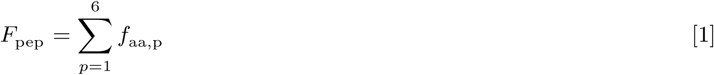

with *f*_aa,p_ the frequency within all self peptides of the AA at position *p* of the 6-mer peptide. These scores were then transformed to a sampling probability *P*_pep_ as follows:

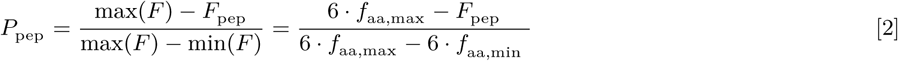

where *f*_aa,max_ is the frequency of the most common AA (L) in all self peptides, and *f*_aa,min_ the frequency of the most rare AA (W). Finally, we sample *n* training peptides from the total set of self peptides using probabilities (*P*_pep_)^s^, where we use the parameter *s* to control the strength of the bias for rare AAs. Throughout the paper, we used either a weak bias (*s* = 1) or a strong bias (*s* = 5) as indicated in the figures.

## Sequence analysis

### String graphs

To visualize strings eliciting reactions from the same TCRs, we constructed a graph where each of 1,000 strings from both languages (English and Xhosa or English and more English) was a node. We then counted for each combination of strings how many TCR motifs (pre-selection) could react to both at *t* = 3, and connected their nodes with an edge if this number was at least 10,000.

For visualization, we ordered the connected components (clusters) in this graph by their number of nodes, and plotted every 10th cluster in the final graph.

### Peptide graphs

To visualize self and foreign peptides to which the same TCRs react, we again started with a graph with nodes for all self-and foreign peptides, and counted for each pair the number of TCRs that could react to both. This time, we used *t* = 4, and connected peptides with an edge if at least 100 TCRs could react to both.

For visualization of HIV and self peptides, we then selected all connected components (clusters) that contained at least one HIV peptide.

### Concordance

Concordances were calculated using the full string-and peptide graphs described above (not just the subsets used for visualization). For each node, we listed the proportion of self-and foreign neighbors. If a node was isolated and had no neighbors, we used the expected value *p*_0,class_ of this proportion (which equals the proportion of self or foreign nodes in the entire graph). For both the self and foreign class of nodes, we then computed the concordance as the mean proportion *p*_class_ of same-class neighbors (so mean proportion of self neighbors for all self nodes, and mean proportion of foreign neighbors for all foreign nodes). Because the ratio between self and foreign peptides/strings was not always equal, we corrected for this ratio as follows:

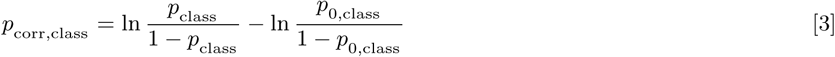

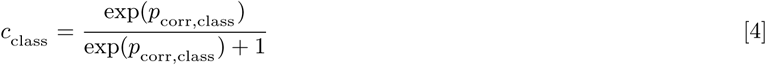

Here, *p*_0,class_ is the expected proportion of same-class neighbors as described above, and *c*_class_ is the ratio-corrected mean concordance for that class (self or foreign). This correction ensures that *c*_class_ = 0.5 when *p*_class_ = *p*_0,class_, 0 when there are only discordant edges between nodes of a different class, and 1 when there are only concordant edges between nodes of the same class. To avoid dividing by zero, we set an exception for situations where *p*_class_ = 1:

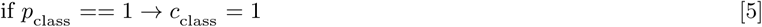

The final, total concordance is then computed as a weighted average of the self-and foreign corrected mean concordance:

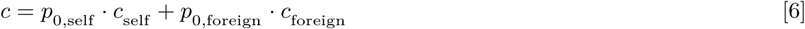

### AA enrichment

The richment of AA *a* (*E*_a_) was computed as

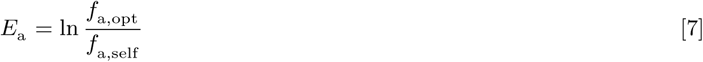

with *f*_a,opt_ the frequency of AA *a* within the optimal set of 130,407 self peptides for *t* = 4 (see *Optimal training peptide selection*), and *f*_a,self_ its frequency within the total set of 263,216 self peptides (Table S1).

### Exchangeability

To compute exchangeability of self peptides, we constructed the graph of all self peptides. We then define exchangeability of a peptide as *N* + 1, where *N* is the number of neighbors in the peptide graph.

To compute how likely peptides of a given exchangeability are to delete foreign-reactive TCRs, we sorted self peptides on their exchangeability and then grouped them into 10 bins with equal numbers of peptides (deciles). Thus, the first decile contains the 10% of peptides with the lowest exchangeabilities, the highest decile the 10% with highest exchangeabilities, etc. We then constructed a graph containing all self and HIV peptides, and analyzed for each decile which percentage of the self peptides in it had an HIV neighbor in this graph (in other words, which percentage “resembled” an HIV peptide).

To analyze the relationship between exchangeability and AA composition, we computed both exchangeability and the AA composition score *F*_pep_ (see *Biased training peptide selection*) for 1000 randomly selected self peptides, and analyzed the association between the two scores.

## Post-selection repertoire analysis

### Sequence recognition

To assess sequence recognition by the post-selection repertoire, we counted the number of post-selection TCRs reacting to each sequence with an affinity of at least the predefined affinity threshold *t* (the same threshold as used for negative selection). Recognition was then reported in the number of reacting TCRs per million TCRs in the post-selection repertoire. If the post-selection repertoire was empty, we set this number to a value of 0. Reported recognition values are always from a single simulation.

### Self-foreign discrimination

To assess self-foreign discrimination within a test set containing equal numbers of self and foreign sequences across multiple simulations, the number of TCRs reacting to each sequence was counted as mentioned above. All sequences were then ranked from high to low numbers of reacting TCRs to obtain the percentage of foreign sequences among the 10% most frequently recognized sequences. When there were ties, we used the value of this percentage that would be expected after random tie-breaking.

### Affinity distribution

To compare TCR affinities between strings to which many TCRs react and strings with fewer reacting TCRs, strings were ranked by number of reacting TCRs as described above and split into the top 10% of most-frequently recognized strings and the remaining 90% of strings. For each string, we then counted the number of TCRs reacting to that string with a specific affinity. For both groups, we then computed how many TCRs recognized a string in that group at a given affinity, and report this as a percentage of all TCRs recognizing a string in that group.

### TCR survival/deletion

To assess TCR survival during negative selection on training sets of increasing size, we first chose a test set of self and/or foreign sequences, and listed all pre-selection TCRs whose affinity for these sequences was ≥ *t*. We then negatively selected our repertoires on training sets that did not contain any of these test sequences, and assessed the percentage of the TCRs of interest that survived negative selection. TCR deletion can then be computed as 100 minus the TCR survival rate.

### Statistical analysis

Central tendency and spread of asymmetrically distributed continuous variables (sequence recognition in TCRs/million) are described using median and interquartile range. For symmetrically distributed continuous variables (% foreign sequences among 10% most frequently recognized sequences, % TCR survival), we use mean and standard deviation (SD). Concordances/AA enrichment scores are computed as a single number for a complete set of sequences and therefore have no measure of spread. The Pearson’s correlation coefficient and 95% confidence interval were computed using the cor.test function of the R stats package with default settings (R version 3.3.2, 2016-10-31, RRID:SCR_001905).

We did not perform frequentist statistical testing, since we can generate as many simulation runs as needed to ensure that any interpreted differences are not simply due to random chance.

**Table S1.** List of proteomes used to extract MHC-I binders. See also *Methods*.

| Organism | Proteome details | Proteins | ID | Download date (d/m/y) | Unique 6-mers (#) |
| --- | --- | --- | --- | --- | --- |
| Ebola virus | Mayinga, Zaire, 1976 | 9 | UP000007209 | 27/09/2017 | 140 |
| Human cyto-megalovirus (HCMV) | Human herpesvirus 5<br>AD169 Isolate<br>Unknown X17403 | 190 | UP000008991 | 27/09/2017 | 2,090 |
| Hepatitis B virus | Genotype D subtype<br>ayw (isolate France/<br>Tiollais/1979) | 7 | UP000007930 | 27/09/2017 | 65 |
| Hepatitis C virus | H77 isolate Unknown<br>AF009606 | 2 | UP000000518 | 27/09/2017 | 112 |
| Human immunodeficiency virus (HIV) | Type 1 group M subtype<br>B (isolate HXB2) | 9 | UP000002241 | 27/09/2017 | 69 |
| Vaccinia virus | Strain Copenhagen | 257 | UP000008269 | 27/09/2017 | 1,955 |
| Zika virus | MR 766 Isolate<br>Unknown AY632535 | 1 | UP000054557 | 27/09/2017 | 118 |
| Listeria monocytogenes | serovar 1/2a<br>(strain ATCC<br>BAA-679 / EGD-e ) | 2,844 | UP000000817 | 27/09/2017 | 31,251 |
| Plasmodium ovale (Malaria) | Wallikeri | 8,636 | UP000078550 | 27/09/2017 | 89,408 |
| Homo sapiens (human) | - | 20,230 | UP000005640 | 01/06/2017 | 263,216 |

**Fig. S1.**
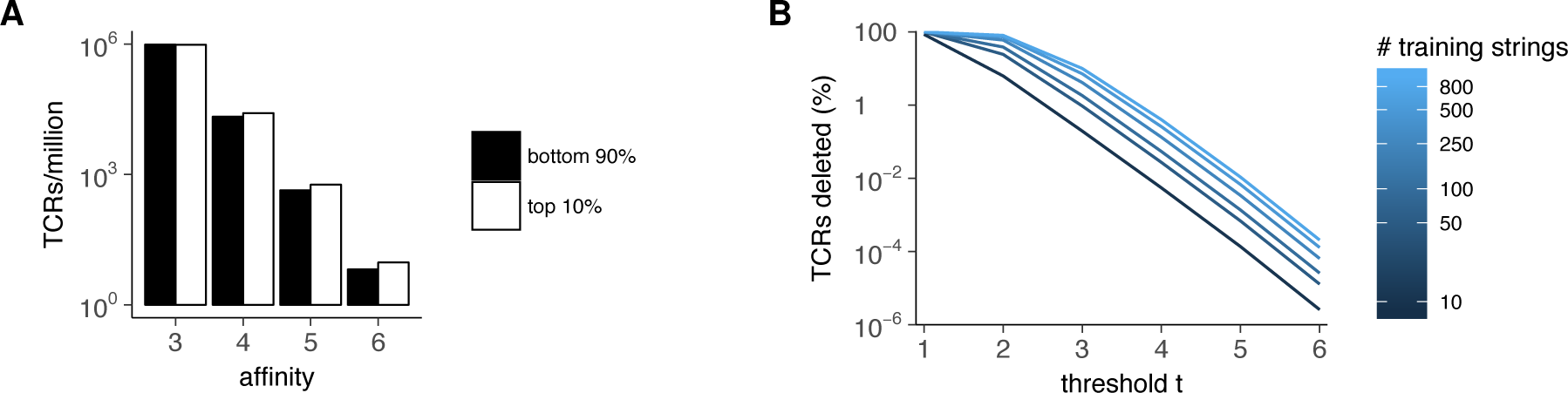
An AIS of string recognition allows simulation of negative selection. (a) Affinity distribution of surviving TCRs reacting to 50 English and 50 Xhosa strings after negative selection. Plot shows TCR counts (of specified affinity) per million total TCRs in either the top 10% of most frequently recognized strings, or the remaining bottom 90% of strings. (b) Average TCR deletion rate as a function of the affinity threshold *t* and the number of training strings used (colored lines). See also Fig. 2A, where we plot these data to show TCR survival as a function of the training set size at *t* = 3.

**Fig. S2.**
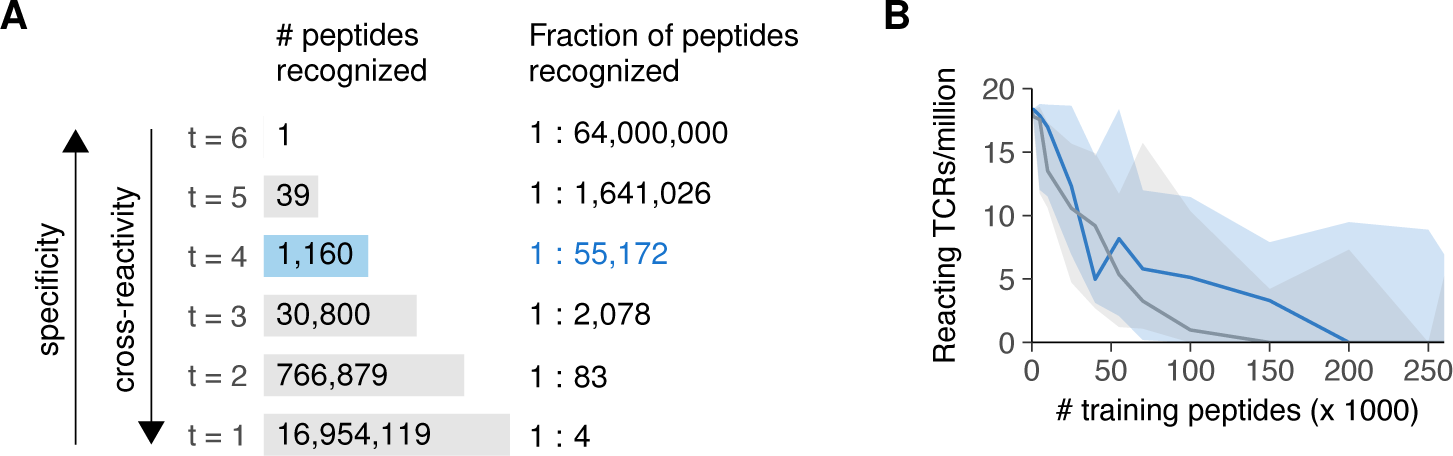
A simple model of TCR-peptide recognition reproduces features of real TCR repertoires. (a) Cross-reactivity at different affinity thresholds *t*. At *t* = 4, a TCR reacts to 1 in every 55,000 peptides, on average. (b) Reanalysis of the data shown in Fig. 3: Typical numbers of TCRs reacting to HIV (blue) and self (gray) peptides after negative selection with *t* = 4. Plot shows median and interquartile range of reacting TCRS/million. Typical values lie between 0 and 20 TCRs per million, depending on the number of training peptides used for negative selection.

**Fig. S3.**
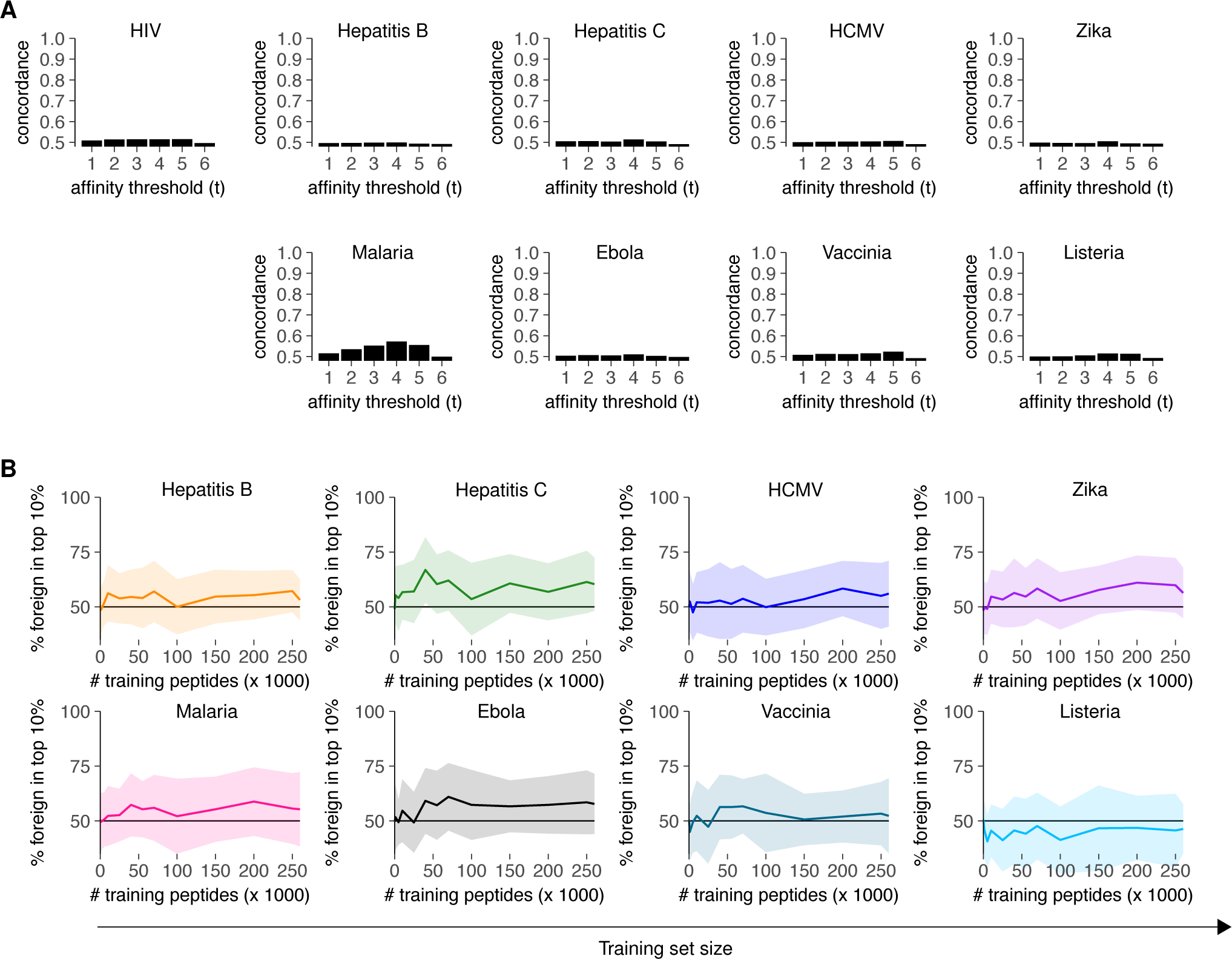
Self-foreign discrimination is poor for all thresholds *t* and all pathogens tested. (a) Concordance (% of same-class neighbors) in the graph of self and foreign peptides is low for all values of *t* and for all pathogens tested. (b) Self-foreign discrimination after negative selection at *t* = 4 is low for all pathogens tested. Plot shows mean*±*SD of the percentage foreign peptides among most frequently recognized peptides (30 simulations).

**Fig. S4.**
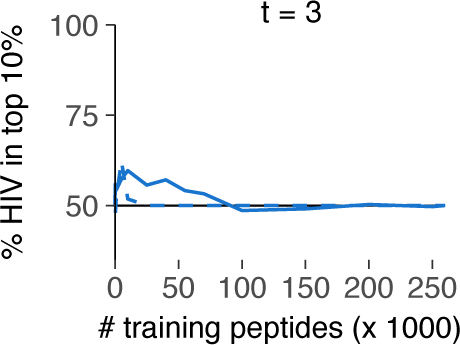
Improved self representation fails to enhance self-foreign discrimination when cross-reactivity is too high. Plot shows mean of the percentage HIV peptides among most frequently recognized peptides after negative selection (*t* = 3, 30 simulations). Negative selection was performed on random (solid line, data from Fig. 3D included for comparison) or optimal (dashed line) training sets.

**Fig. S5.**
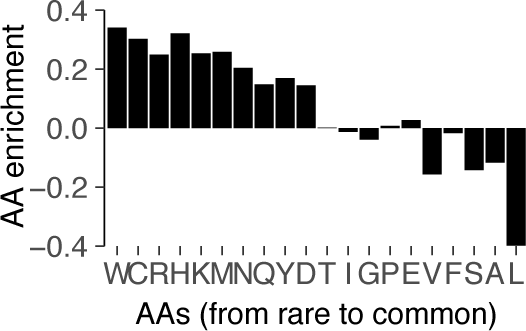
Optimal training sets are enriched for rare AAs. Plot shows AA enrichment in optimal training set. Enrichment is the log of the observed frequency divided by the frequency among all self peptides. Negative values indicate depletion.

**Fig. S6.**
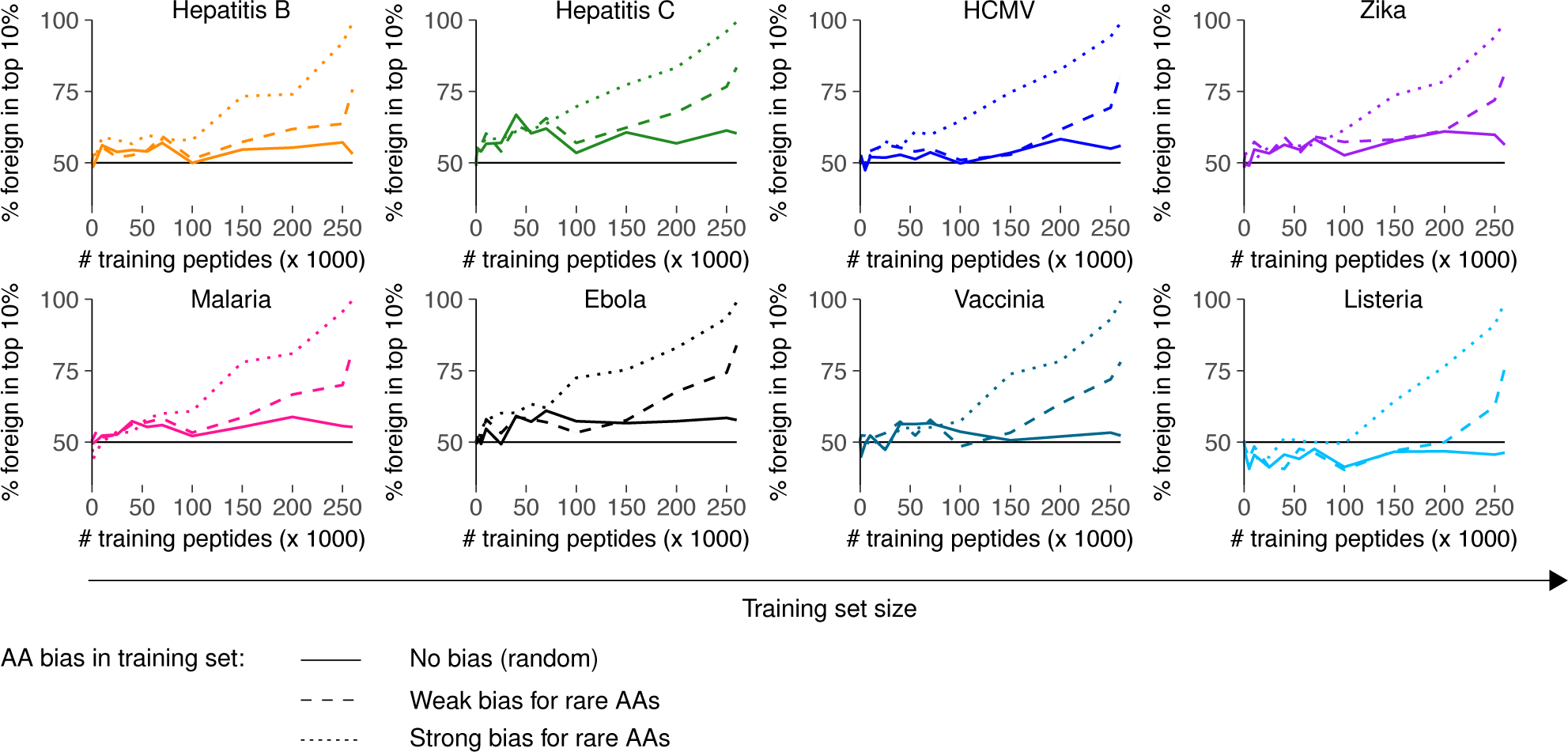
Increased presentation of rare AAs during negative selection improves self-foreign discrimination for all pathogens tested. Plot shows mean±SD of the percentage foreign peptides among most frequently recognized peptides after negative selection (*t* = 4, 30 simulations). Training peptides were either chosen randomly (solid line, data from Fig. S3B included for comparison) or with a weak/strong bias for peptides with rare AAs (dashed/dotted lines).

